# Role of maternal, fetal and placental histopathology factors in the pathogenesis of retinopathy of prematurity

**DOI:** 10.1101/2023.02.13.528236

**Authors:** Tarandeep Kaur, Neha Sharma, Saumya Jakati, Nitasha Bagga, Sanchita Mitra, Katamtreddy Bhargavi K, Subhadra Jalali, Inderjeet Kaur

**Affiliations:** Kallam Anji Reddy Molecular Genetics Lab, Brien Holder Eye Research Centre, Hyderabad Eye Research Foundation, LV Prasad Eye Institute, KAR Campus, Hyderabad, India; Ophthalmic pathology laboratory, LV Prasad Eye Institute, KAR Campus, Hyderabad, India; Department of Neonatology, Rainbow Children’s Hospital, Banjara Hills, Hyderabad, India; Jhaveri Microbiology Centre & Saroja A Rao Immunology Laboratory, LV Prasad Eye Institute, KAR Campus, Hyderabad, India; Department of Obstetrics and Gynecology, Rainbow Children’s Hospital, Banjara Hills, Hyderabad, India; Department of Vitreo-retina, LV Prasad Eye Institute, KAR Campus, Hyderabad, India

## Abstract

**Purpose:** To investigate if maternal, fetal and placental vascular and/or molecular changes predict the risk for retinopathy of prematurity in preterm infants.

**Methodology:** The postnatal and antenatal data and placental histopathological changes; H&E staining of placental sections and molecular findings; gene expression analysis by qRT-PCR and protein expression by IHC at the maternal-fetal interface were collected from 20 placental samples and categorized into 3 groups: full-term (n=10), preterm without ROP (n=7) and preterm with ROP (n=6).

**Results:** The correlation analysis indicated significant association of increased monocytes (p=0.042), fetal growth retardation (p=0.000), apnoeic spell (p=0.033), ventilation (p=0.009), length in hospital stay (p=0.001) and decreased RBC (p=0.02), Hb (p=0.048), PCV (p=0.010), gestational age (p=0.003), birth weight (p=0.000) with increased risk of ROP. Along with these factors, placental weight(p=0.001), diameter (p=0.019), Tenny-Parker changes (p=0.025), alternating area of small and short hyper mature villi (p=0.033) are also found to be significant determinants of the disease. Gene expression analysis revealed significant increase in hypoxia (*HIF-1* gene expression; p=0.007) and non-significant increase in the pro-inflammatory marker *IL-6*. The protein expression also showed the significantly increased activation of complement pathway (CFH) and NF- ΚB at maternal-fetal interface.

**Conclusions:** Our preliminary results support the changes in the maternal-fetal factors, placental histopathology and molecular alterations related to hypoxia, inflammation and complement activation at maternal-fetal interface in preterm with ROP placentas. These changes have the potential to predict the risk of the disease but the results are required to be further validated in larger cohort.

## Introduction

Retinopathy of prematurity (ROP) is a potentially blinding vaso-proliferative condition of retina of the eye, seen predominantly, in premature babies. Despite the advancement of childcare modalities, the preterm infants due to incomplete or delayed development can sometimes face permanent learning, hearing and visual disabilities (Menon *et al*., 2012). Retinopathy of prematurity (ROP) can contribute to visual impairment or blindness, especially if not screened and treated on time. The incidence of ROP varies from 19.88% (Ludwig *et al*., 2017) in developed countries with well-equipped neonatal intensive care units to ≈ 30% (Gergely and Gerinec, 2010) in developing countries. In India, the incidence varies from 38-51.9% in different states and out of 26 million low birth babies, 2 million babies are at risk of developing ROP annually (Pejawar *et al*., 2010) Low birth weight and gestational age are considered as the most important risk factors for the development of ROP in developed countries. Besides these large number of other risk factors including sepsis, vitamin E deficiency, multiple births, blood transfusion, oxygen supplementation, ethnicity, prolonged mechanical ventilation, hyperglycemia, anemia, diabetes, pre-eclampsia and many more factors, cumulatively impact the emergence of ROP (Alajbegovic-Halimic *et al*., 2015). However, some preterm infants are seen to develop severe, vision threatening ROP even in the absence of any of these known risk factors; on the opposite spectrum are babies who show spontaneous regression of ROP. Therefore, it becomes important to identify the preterm infants that are at increased risk of developing severe ROP. Secondly, ROP must be detected as early as possible to initiate immediate treatment because present treatment options including laser ablation, anti-VEGF injections and vitrectomy surgery can only stop the progression of the disease; the anatomical changes that have already occurred cannot be reversed and many children show unsatisfactory visual outcomes (Holmström and Larsson, 2008). Therefore, there is a need of early diagnosis/predictive testing for ROP, so that the disease course can be modified much before the clinical features of abnormal vessel proliferation are seen.

Many clinical and experimental studies have shown that early life conditions powerfully influence the later life susceptibility to diseases. Even in case of ROP, it has been reported that chorioamnionitis and placental ischemia are associated with increased risk of ROP due to induction of inflammatory mechanisms (Mitra *et al*., 2014) or due to decrease in the IGF-1 levels because of inflammation (Moscuzza *et al*., 2011). Along with the inflammatory factors few reports have shown the association of cord blood plasma IL-6 and C5a protein levels to be significantly associated with severe ROP and laser treatment respectively (Park *et al*., 2019). These studies predict the involvement of prenatal/in-utero pathological events that can lead to ROP in postnatal life. But all these reports are isolated reports without any validation or detailed investigation and most of the studies on ROP till now are done after the birth of the preterm babies.

One of the vital organs in utero which maintain balance between maternal and fetal environment during pregnancy is the placenta. Functionally, it is a complex organ that performs circulatory, hormonal and immunological functions of the fetus and the mother (Simister, 2003.) and synthesizes several molecules essential for normal development of placenta itself, metabolism of mother, normal fetal growth and parturition. Even the expression of large number of genes involved in normal succession of pregnancy is regulated by growth factors and hormones secreted by placenta. All these processes ultimately depend upon the normal vascular development of placenta whereas insufficient/abnormal placental vasculature can result into complications like preeclampsia, fetal growth retardation (Wang *et al*., 2010) and even incomplete vasculature formation in the retina that are directly associated with future programming of complications like ROP.

The complement cascade is another factor that plays an important role in placental and fetal development and proper complement system activation is required from pre-implantation to parturition and to avoid adverse pregnancy outcomes (Girardi *et al*., 2020). The complement cascade also plays an important role in the normal neuron and vasculature formation. Thus, compromised neurovascular development during pregnancy due to altered levels of complement activation in the retina can predispose preterm infants to severe ROP. In a previous study from our lab, we have found strong association of variants in complement factor H and complement factors B genes with ROP. Along with these genes proteins involved in complement pathway were also significantly increased in the extracellular matrix and vitreous of ROP babies (Rathi *et al*., 2017). This abnormal immune complement components in the eyes of ROP babies can be a consequence of *in-utero* changes or postnatal effects and needs further study.

Keeping in view the importance of placenta, inflammation and complement activation during pregnancy, the analysis of structure, gene expression profiling of complement system related markers and the inflammatory markers in placental tissue becomes important along with the antenatal and postnatal maternal and fetal factors when evaluating the pathogenesis of ROP.

## Materials and methods

### Study design

A case-control study approach was adopted. The prospective study adhered to the tenets of the declaration of Helsinki. The maternal and infant demographical and clinical features along with placental vascular lesions and molecular changes at maternal-fetal interface were compared between full term, preterm without ROP and preterm with ROP groups. The study was carried out under written informed consent from all the participants and was approved by Institutional Review Board (LEC-BHR-01-20-380) of the LV Prasad Eye Institute (LVPEI).

### Study participants

After written informed consent either from the father or the mother, placentas from 46 lower (uterine) segment caesarean section (LSCS) deliveries at Rainbow children’s Hospital, Hyderabad, India were collected for the study. The detailed prenatal and antenatal information of the mother and infant was also recorded on a predesigned questionnaire. The placentas from spontaneous conceptions and singleton deliveries with gestational age ≤34 weeks and birth weight ≤1700g were included in the preterm study group while the deliveries with GA≥36 weeks and birth weight ≥1700g were included in full-term group. The placentas from assisted reproductive technology (ARTs) conceptions, pregnancies complicated by any type of infections, congenital and/or chromosomal disorders, family history of any other retinal disorders and intraocular infections were excluded from the study. All the infants underwent detailed clinical examination by a neonatologist and preterm infants with intrauterine growth retardation were also excluded from the study. The included preterm infants were thoroughly examined by an ophthalmologist at 21^st^ day of life for retinal vascular status as part of the routine screening done by the retina specialists of LVPEI. maturation. Based upon the examination the preterm infants were further divided into preterm without ROP and preterm with ROP group.

### Placental sample collection and processing

All the placental samples (full-term=10; pre-term without ROP= 07 and preterm with ROP=06) were collected within half an hour of delivery and were transported to LVPEI on ice in aseptic conditions. The diameter and weight of each fresh placenta sample was recorded on a specialized performa. The weight of the placenta was measured after trimming the umbilical cord as per the Amsterdam placental workshop group consensus statement (Khong *et al*., 2016). The placental samples were washed with chilled 1X PBS, grossed on ice to take 4-6 specific sections of decidua and chorionic villi for histopathological and gene expression analysis (from the same areas). More sections were grossed from the site of any visible pathological change.

### Microbiological investigations

To rule out the role of infections in ROP, small section of the tissues were used for culture based and PCR based microbiological investigations for bacteria, fungus and viral infections. A small tissue chunk of each collected placenta was directly analysed for culture based on blood agar and brain heart infusion media growth. For PCR based investigations the primer details for bacteria, fungus and virus are given in table 1.

**Table 1:** Maternal, fetal and placental continuous variables in three different groups:

| Factor | Full term | Preterm | Preterm with ROP | full term VS Preterm (student's t-test p-value) | preterm VS Preterm-ROP (student's t-test p-value) |
| --- | --- | --- | --- | --- | --- |
| Maternal Age | 30.57±1.78 | 31.43±1.94 | 34±1.12 | .751 | .279 |
| Hb (g/Dl) (12-14) | 11.77±0.68 | 12.03±0.36 | 11.12±0.15 | .747 | **.046** |
| red blood cells (milli/mm<sup>3</sup>) (4-5.5) | 4.37±0.19 | 4.27±0.09 | 3.87±0.11 | .648 | **.021** |
| PACKED CELL VOLUME (PCV) (volume%) (37-47) | 36.04±1.74 | 36.26±0.77 | 33.27±0.55 | .913 | **.009** |
| Mean Corpuscular Volume (MCV) (fL) (80-99) | 82.43±1.29 | 85.09±1.56 | 84.52±1.87 | .216 | .820 |
| MCH (pg/cells) (23-31) | 26.85±0.78 | 28.24±0.86 | 28.3±0.59 | .258 | .957 |
| MCHC (g/dl) (33-37) | 32.57±0.57 | 33.19±0.51 | 33.47±0.15 | .439 | .613 |
| PLATELET COUNT (Lakhs/mm<sup>3</sup>) (1.5-4.5) | 2.94±0.20 | 2.002±0.15 | 2.38±0.24 | **.003** | .221 |
| WBC COUNT (Cells/mm<sup>3</sup>) (4500-10500) | 10795.71±1050.50 | 13714.29±1645.73 | 16051.67±1740.75 | .165 | .351 |
| MPV (fL) (8-12) | 10.07±0.35 | 8.96±1.53 | 10.13±0.32 | .501 | .477 |
| NUETROPHILS (50-70) | 75±3.17 | 73.86±3.12 | 77.33±3.25 | .802 | .457 |
| LYMPHOCYTES (30-50) | 20.14±2.95 | 21.43±2.94 | 18.67±4.22 | .763 | .604 |
| MONOCYTES (1-10) | 2.14±0.26 | 2.29±0.29 | 3.67±0.56 | .718 | .061 |
| EOSINOPHILS (1-6) | 2.71±0.29 | 2.43±0.20 | 2±0.26 | .432 | .221 |
| RDW | 15.26±0.67 | 14.16±0.67 | 14.65±0.50 | .270 | .569 |
| SBP | 119.29±4.14 | 137±6.86 | 140.5±8.05 | **.052** | .747 |
| DBP | 76.43±2.83 | 93.29±3.88 | 89.17±7.35 | **.005** | .634 |
| Gestational Age | 36.71±0.61 | 34.43±0.29 | 29.83±1.25 | **0.008** | **.013** |
| Birth weight | 2.57±0.14 | 2.32±0.13 | 1.25±0.10 | .199 | **.000** |
| Placental weight | 0.49±0.07 | 0.44±0.03 | 0.29±0.02 | 0.610 | **.001** |
| Placental diameter | 20.93±0.60 | 18.07±0.98 | 14.6±0.75 | **.032** | **.017** |
| Apgar Score at 1 min | 8.71±0.18 | 8±0.58 | 7.5±0.22 | .276 | .443 |
| Apgar score at 5 min | 8.86±0.26 | 9±0.22 | 8.33±0.33 | .682 | .129 |
| Length of hospital stay | 2.14±0.70 | 3.29±0.52 | 27±5.85 | .032 | **.010** |

### Histopathology

All the placentas were examined for gross and microscopic features. Placentas were fixed overnight in 10% neutral buffer formalin. Once fixed, the placenta was sub-divided into umbilical cord, membranes, the parenchyma and were assessed for parameters including shape, size, colour and morphological abnormalities. Tissue sections were taken from central middle and peripheral protocol regions and the areas with abnormalities. Tissue sections were processed in automated tissue processor (Leica TP 1020) and embedded in liquid paraffin to create tissue blocks. Blocks were sectioned at 4-micron thickness on a microtome (Fisher *et al*., 2008a) and stained with Haematoxylin and Eosin (H&E) stain. (Fisher *et al*., 2008b). The microscopy slides were scanned under Leica APERIO AT2 digital pathology scanner and examined by a pathologist blinded to the ROP status of the infants.

### RNA extraction, cDNA conversion and semi-quantitative PCR

The RNA was extracted from each tissue section by minor modifications of the Trizol extraction method (Rio *et al*., 2010). The quality and quantity of the RNA samples were analysed by UV-absorbance spectrophotometry and integrity was checked using a 2100 Bioanalyzer platform (Agilent Technologies). The cDNA was prepared from extracted RNA samples using iScript cDNA synthesis kit (Bio-Rad, CA, USA). The prepared cDNA was then used for microbiological investigations by conventional PCR followed by agarose gel electrophoresis. The semi quantitative PCR was performed to evaluate the gene expression for oxidative stress (*HIF1*, *NRF2*), complement system (*C3*, *CFH*), Inflammatory (*IL6*, TNFα) and anti-inflammatory (*IL4*, *IL10*) markers.

### Immunohistochemistry for validation of gene expression

The tissue sections were deparaffinized using xylene followed by gradual rehydration with ethanol. The antigen retrieval was performed using microwave technique wherein the tissue section is placed in sodium citrate buffer and heated twice at full power for 4-5 minutes each followed by a single cycle at medium power for 4-5 minutes. After antigen retrieval permeabilization was achieved by incubation of tissue sections with 5% triton-x 100. The blocking of non-specific antibodies was performed by incubation with 2% BSA in phosphate buffer saline (PBS) for 45 minutes at room temperature. Thereafter the sections were incubated overnight at 4°C with primary antibody (the primary antibodies for C3 and CFH were monoclonal from Santa Cruz raised in mouse and for NF-KB the monoclonal primary antibody was from Biolegend raised in mouse) diluted in 1X PBS followed by washing with 1X PBS and incubation for 60 minutes with biotinylated anti-mouse immunoglobulin. The sections were then stained with DAPI for nuclei staining. The slides were then mounted and scanned under EVOS microscope. Images were captured and scanned by ImageJ software to quantify the fluorescence signal (Schindelin *et al*., 2015).

### Statistical analysis

The placental weight and diameter are represented as mean±SE and compared in different groups by student’s t-test. The p value ≤0.05 was considered as statistically significant. The correlation analysis followed by multiple linear regression analysis was used to find out the association if any between the prenatal and antenatal maternal and fetal factors and placental histopathological features with the disease status. The gene expression and fluorescence quantification data were represented as scatter plots and was compared by student’s t-test in different groups.

## Results

### Study participants

In total 39 placenta samples were collected for the study and categorized into full term (n=29), preterm with no ROP (n=7) and preterm that were diagnosed with different stages of ROP (n=6). The maternal and fetal clinical characteristics of all three groups and histopathological evaluation of placental samples (full term=10; preterm, with no ROP=7; preterm with ROP=6) are represented in Table: 1.

### Microbiological investigations of placenta samples

The culture based microbiological investigations showed the infection of *Pseudomonas* and *Ralstonia* sp. in 07 of the 29 full-term placental samples. To remove the confounding role of these infections, these 07 placental samples were excluded from the study. PCR based investigations showed that all the other placental samples were bacteria, fungus and HSV viruses infection free.

### Maternal and fetal variables

There was significant difference of maternal platelet count (p=0.003), systolic blood pressure (SBP; p=0.052), diastolic blood pressure (DBP; p=0.005), gestational age (p=0.008) and placental diameter (p=0.032) among full term and preterm group individuals. When these factors were compared between preterms with no ROP and preterm with ROP groups, maternal haemoglobin (Hb), red blood cell count (RBC), packed cell volume (PCV) were significantly different among the two groups in the third trimester of pregnancy (p values= 0.046, 0.021, 0.009, respectively). Along with these maternal factors, gestational age (29.83±1.25weeks; p= 0.013), birth weight (1.25±0.10kg; p=0.000), placental weight (0.29±0.02kb; p=0.001) and placental diameter (14.6±0.75cm; p=0.017) were significantly decreased while length of hospital stay was significantly higher (27±5.85days; p=0.010) in preterm with ROP group individuals compared to preterm without ROP.

In preterm with ROP cases, maternal preeclampsia/gestational hypertension and fetal distress during pregnancy was present in 83% of the mothers while in preterm with no ROP and full-term infants, it was 71% and 29% cases, respectively (Table: 2). After birth, 83% of infants were having respiratory distress and given ventilation support and that were diagnosed with ROP later. The respiratory distress was present only in 71% preterm infants with no-ROP and only 14% were given ventilation support.

**Table 2:** Distribution of maternal, fetal and placental categorical variables in three different groups:

|  | Full-term<br>Placentae<br>(n=7)<br>(present/absent) | Preterm<br>without ROP<br>(n=7)<br>(present/absent) | Preterm with<br>ROP<br>(n=6)<br>(present/absent) |
| --- | --- | --- | --- |
| **Maternal factors** |  |  |  |
| Assisted conception | 1/6 (14%) | 1/6 (14%) | 1/5 (17%) |
| Pre-eclampsia/gestational hypertension | - | 5/2 (71%) | 5/1 (83%) |
| PPROM | 1/6 (14%) | 1/6 (14%) | 2/4 (33%) |
| Amniotic fluid abnormalities | - | 1/6 (14%) | - |
| Fetal distress | 1/6 (14%) | 2/5 (29%) | 5/1 (83%) |
| Placental abruption | - | - | 1/5 (17%) |
| Diabetes | - | 3/4 (43%) | 2/4 (33%) |
| Thyroid | 1/6 (14%) | 1/6 (14%) | 2/4 (33%) |
| **Fetal factors** |  |  |  |
| Fetal growth Retardation | - | - | 6/0 (33%) |
| Respiratory distress | 1/6 (14%) | 5/2 (71%) | 5/1 (83%) |
| Pneumothorax | - | 1/6 (14%) | - |
| Bronchitis | - | 1/6 (14%) | - |
| Apnoea | - | 1/6 (14%) | 3/3 (50%) |
| Apnoeic spell | - | - | 3/3 (50%) |
| Brady spell | - | 1/6 (14%) | - |
| Bacterial/fungal sepsis | - | 1/6 (14%) | 1/5 (17%) |
| Anaemia | - | - | 1/5 (17%) |
| Hyperglycaemia | 1/6 (14%) | - | 1/5 (17%) |
| Oxygen | - | - | 1/5 (17%) |
| Surfactant | - | - | 1/5 (17%) |
| Ventilation | 1/6 (14%) | 1/6 (14%) | 5/1 (83%) |
| Hyperbilirubinemia | 1/6 (14%) | 5/2 (71%) | 3/3 (50%) |
| Phototherapy | 2/5 (29%) | 4/3 (57%) | 3/3 (50%) |
| **Placental factors** |  |  |  |
| Arterial thrombosis | - | - | 1/5 (17%) |
| Distal villous hypoplasia | - | - | 2/4 (33%) |
| Thin elongated villi | 2/5 (29%) | 4/3 (57%) | 6/0 (100%) |
| Tenny Parker changes | 3/4 (43%) | 3/4 (43%) | 6/0 (100%) |
| Alternating area of small or short hyper mature villi | - | - | 3/3 (50%) |
| Agglutinated villi |  |  | 3/3 (50%) |
| Increased syncytial knots | 5/2 (71%) | 7/0 (100%) | 6/0 (100%) |
| Intervillous fibrin | - | - | 2/4 (33%) |
| Villous paucity due to decreased villous branching | - | - | 1/5 (17%) |
| Crowding and congestion of villi | 3/4 (43%) | 4/3 (57%) | 2/4 (33%) |
| Loss of nuclear staining in stroma | - | - | 2/4 (33%) |
| Ghost villi | - | - | 2/4 (33%) |
| Loss of trophoblastic villi | - | - | 2/4 (33%) |
| Perivillous fibrin deposition | - | 2/5 (29%) | 2/4 (33%) |
| Intervillous haemorrhage | 1/6 (14%) | 1/6 (14%) | 2/4 (33%) |
| Delayed villous maturation [Decreased blood vessels in terminal villi] | - | - | 1/5 (17%) |
| Chorangiosis (hyper capillarization of terminal villi) | - | - | 1/5 (17%) |
| Multiple chorangiomatosis (involving small vessels at periphery of immature intermediate villi) | - | - | 1/5 (17%) |
| Intramural fibrin in large fetoplacental vessels | - | - | 1/5 (17%) |
| Chorioamnionitis | - | - | 1/5 (17%) |
| Subchorionitis | - | - | 1/5 (17%) |
| Intervillositis and perivillous fibrin deposition (Affects intervillous space) | - | 1/6 (14%) | - |
| Abnormal insertion | - | 1/6 (14%) | 2/4 (33%) |

### Association of maternal and fetal variables with ROP

The Person’s correlation followed by multiple linear regression analysis was performed after categorizing the data into three groups (full term, preterm with no ROP and preterm with ROP) and into two groups (preterm with no ROP and preterm with ROP) in order to identify variables that are associated with the disease risk.

The determinants of the disease when study group is categorized into three category were RBC (p=0.029), WBC (p=0.021), monocytes (p=0.014), SBP (p=0.03), preeclampsia (p=0.001), fetal distress (p=0.011), gestational age (p=0.000), birth weight (p=0.000), APGAR at 1min (p=0.036), fetal growth retardation (p=0.000), respiratory distress (p=0.009), apnoea (p=0.026), apnoeic spell (p=0.012), ventilation (p=0.009), nutrition (p=0.011) and length of hospital stay (p=0.000). Among the associated variables decreased RBC count in the third trimester, lower gestational age and birth weight are at increased risk for the disease while human milk is protective for the disease.

When correlation analysis was performed after categorising into two groups i.e., preterm with no ROP and preterm with ROP, increased monocytes in third trimester (p=0.042), fetal growth retardation (p=0.000), apnoeic spell (p=0.033), ventilation (p=0.009) and length of hospital stay (p=0.001) are significantly associated with the disease while decreased RBC (p=0.02), Hb (p=0.048), PCV (p=0.010), gestational age (p=0.003) and birth weight (p=0.000) increases the risk of the disease. But after multiple linear regression analysis only fetal growth retardation was significantly associated with ROP (Table: 3).

**Table 3:** Association of maternal and fetal factors with the disease state.

|  |  | Full term,<br>preterm, preterm<br>with ROP | preterm with ROP<br>and preterm<br>without ROP<br>groups |
| --- | --- | --- | --- |
| Maternal Age | Pearson Correlation | 0.316 | 0.314 |
|  | Sig. (2-tailed) | 0.174 | 0.297 |
| Hb | Pearson Correlation | -0.210 | **-.557\*** |
|  | Sig. (2-tailed) | 0.374 | **0.048** |
| RBC | Pearson Correlation | **-.489\*** | **-.633\*** |
|  | Sig. (2-tailed) | **0.029** | **0.020** |
| PCV | Pearson Correlation | -0.350 | **-.684\*\*** |
|  | Sig. (2-tailed) | 0.130 | **0.010** |
| MCV | Pearson Correlation | 0.224 | -0.071 |
|  | Sig. (2-tailed) | 0.343 | 0.818 |
| MCH | Pearson Correlation | 0.304 | 0.016 |
|  | Sig. (2-tailed) | 0.192 | 0.959 |
| MCHC | Pearson Correlation | 0.307 | 0.147 |
|  | Sig. (2-tailed) | 0.188 | 0.633 |
| Platelet count | Pearson Correlation | -0.395 | 0.379 |
|  | Sig. (2-tailed) | 0.085 | 0.202 |
| WBC count | Pearson Correlation | **.513\*** | 0.282 |
|  | Sig. (2-tailed) | **0.021** | 0.351 |
| MPV | Pearson Correlation | 0.000 | 0.206 |
|  | Sig. (2-tailed) | 1.000 | 0.500 |
| Neutrophils | Pearson Correlation | 0.115 | 0.226 |
|  | Sig. (2-tailed) | 0.628 | 0.458 |
| lymphocytes | Pearson Correlation | -0.069 | -0.164 |
|  | Sig. (2-tailed) | 0.774 | 0.593 |
| monocytes | Pearson Correlation | **.541\*** | **.571\*** |
|  | Sig. (2-tailed) | **0.014** | **0.042** |
| Eosinophils | Pearson Correlation | -0.431 | -0.371 |
|  | Sig. (2-tailed) | 0.058 | 0.212 |
| RDW | Pearson Correlation | -0.166 | 0.170 |
|  | Sig. (2-tailed) | 0.485 | 0.579 |
| SBP | Pearson Correlation | **.486\*** | 0.100 |
|  | Sig. (2-tailed) | **0.030** | 0.745 |
| DBP | Pearson Correlation | 0.396 | -0.154 |
|  | Sig. (2-tailed) | 0.084 | 0.615 |
| ART | Pearson Correlation | 0.026 | 0.033 |
|  | Sig. (2-tailed) | 0.913 | 0.915 |
| Preeclampsia | Pearson Correlation | **.684\*\*** | 0.141 |
|  | Sig. (2-tailed) | **0.001** | 0.646 |
| Amniotic fluid anomalies | Pearson Correlation | 0.014 | -0.267 |
|  | Sig. (2-tailed) | 0.952 | 0.377 |
| Fetal distress | Pearson Correlation | **.558\*** | 0.548 |
|  | Sig. (2-tailed) | **0.011** | 0.053 |
| Placental abruption | Pearson Correlation | 0.299 | 0.312 |
|  | Sig. (2-tailed) | 0.200 | 0.300 |
| Thyroid | Pearson Correlation | 0.186 | 0.225 |
|  | Sig. (2-tailed) | 0.431 | 0.459 |
| Gestational age | Pearson Correlation | **-.822\*\*** | **-.758\*\*** |
|  | Sig. (2-tailed) | **0.000** | **0.003** |
| Birth weight | Pearson Correlation | **-.826\*\*** | **-.889\*\*** |
|  | Sig. (2-tailed) | **0.000** | **0.000** |
| Apar 1min | Pearson Correlation | **-.470\*** | -0.223 |
|  | Sig. (2-tailed) | **0.036** | 0.465 |
| Apgar 5min | Pearson Correlation | -0.289 | -0.461 |
|  | Sig. (2-tailed) | 0.216 | 0.113 |
| FGR | Pearson Correlation | **.854\*\*** | **1.000\*\*** |
|  | Sig. (2-tailed) | **0.000** | **0.000** |
| Respiratory distress | Pearson Correlation | **.568\*\*** | 0.141 |
|  | Sig. (2-tailed) | **0.009** | 0.646 |
| Pneumothorex | Pearson Correlation | 0.014 | -0.267 |
|  | Sig. (2-tailed) | 0.952 | 0.377 |
| Bronchitis | Pearson Correlation | 0.014 | -0.267 |
|  | Sig. (2-tailed) | 0.952 | 0.377 |
| Aponea | Pearson Correlation | **.497\*** | 0.386 |
|  | Sig. (2-tailed) | **0.026** | 0.193 |
| Apnoeic spell | Pearson Correlation | **.548\*** | **.592\*** |
|  | Sig. (2-tailed) | **0.012** | **0.033** |
| Brady spell | Pearson Correlation | 0.014 | -0.267 |
|  | Sig. (2-tailed) | 0.952 | 0.377 |
| Bacterial fungal sepsis | Pearson Correlation | 0.228 | 0.033 |
|  | Sig. (2-tailed) | 0.334 | 0.915 |
| Anaemia | Pearson Correlation | 0.299 | 0.312 |
|  | Sig. (2-tailed) | 0.200 | 0.300 |
| Hyperglycemia | Pearson Correlation | 0.021 | 0.312 |
|  | Sig. (2-tailed) | 0.931 | 0.300 |
| Oxygen | Pearson Correlation | 0.228 | 0.033 |
|  | Sig. (2-tailed) | 0.334 | 0.915 |
| No of days oxygen | Pearson Correlation | 0.150 | -0.103 |
|  | Sig. (2-tailed) | 0.528 | 0.738 |
| Surfactant | Pearson Correlation | 0.299 | 0.312 |
|  | Sig. (2-tailed) | 0.200 | 0.300 |
| Ventilation | Pearson Correlation | **.567\*\*** | **.690\*\*** |
|  | Sig. (2-tailed) | **0.009** | **0.009** |
| Hyperbilirubenia | Pearson Correlation | 0.306 | -0.220 |
|  | Sig. (2-tailed) | 0.189 | 0.471 |
| Phototherapy | Pearson Correlation | 0.181 | -0.071 |
|  | Sig. (2-tailed) | 0.445 | 0.817 |
| Phototherapy duration | Pearson Correlation | -0.039 | -0.263 |
|  | Sig. (2-tailed) | 0.870 | 0.385 |
| Nutrition | Pearson Correlation | **-.556\*** | **-.857\*\*** |
|  | Sig. (2-tailed) | **0.011** | **0.000** |
| Length of hospital stay | Pearson Correlation | **.733\*\*** | **.798\*\*** |
|  | Sig. (2-tailed) | **0.000** | **0.001** |

### Placental histopathological changes at maternal-fetal interface in ROP

The placental continuous variables are presented in Table: 1, while categorical variables are summarized in Table: 2. The average placental diameter was significantly decreased in preterm with ROP group individuals (14.6±0.75cm) compared to preterm with no ROP (18.07±0.98cm) and full term (20.93±0.60cm) with p-value equals to 0.017 and 0.032, respectively. Along with the placental diameter, placental weight was also significantly lesser in preterm with ROP group (0.29±0.02kg; p=0.001). The maternal vascular lesion i.e., arterial thrombosis was present in 15% of placentas from preterm deliveries and infants that developed ROP. Features of maternal malperfusion including distal villous hypoplasia, alternating areas of small or short hyper mature villi, agglutinated villi, intervillous fibrin, villous paucity due to decreased villous branching and loss of nuclear staining were observed only in 17%-33% of placenta samples form preterm with ROP category (Table: 2). Besides these maternal placental lesions few fetal vascular lesions including delayed villous maturation, chorangiosis and multiple chorangiomatosis have only been observed in the placentas from preterm with ROP cases.

### Association of placental histopathological variables with the disease risk

After stratification of studied samples into three groups, correlation analysis revealed placental weight, diameter, distal villous hypoplasia, thin elongated villi, Tenny - Parker changes, alternating area of small and short hyper mature villi to be significant determinants of the disease. As the samples were further stratified into two groups preterm without and preterm with ROP, parameters such as distal villous hypoplasia and thin elongated villi lost the significance (Table 4).

**Table 4:**
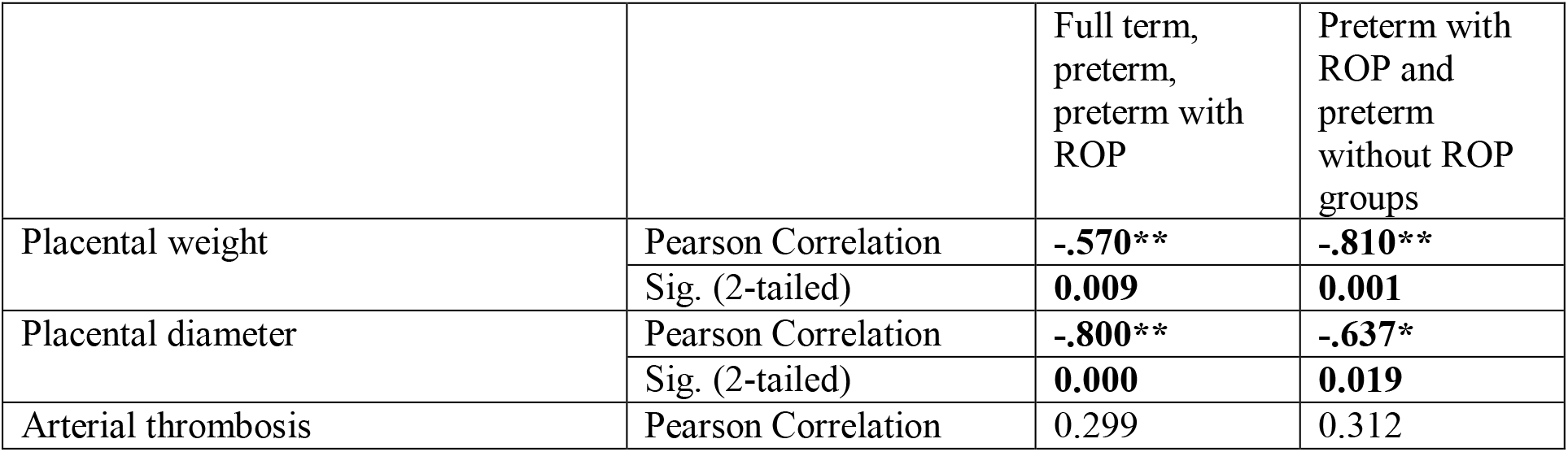
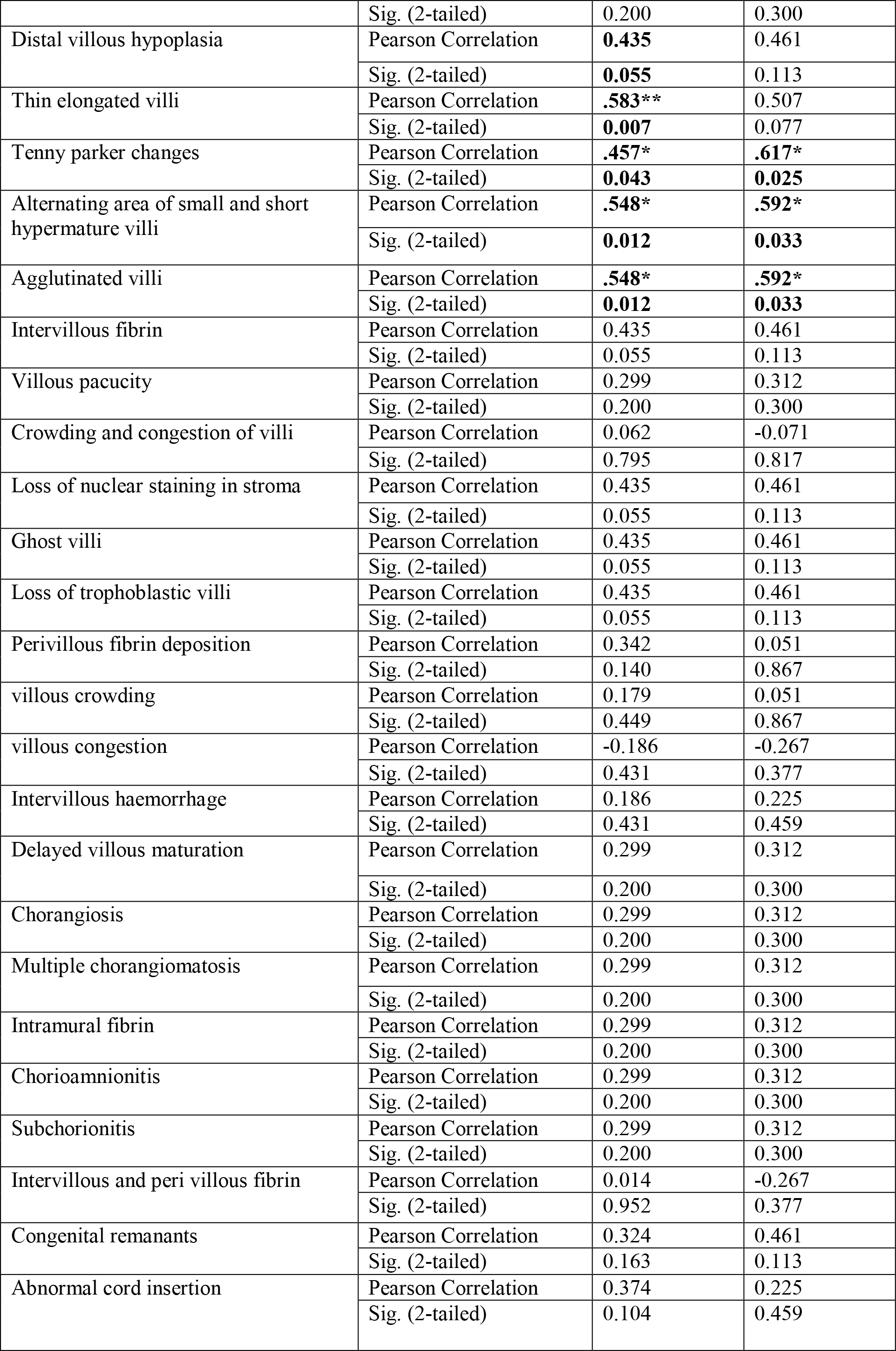
Association of placental factors with the disease state.

The observed histopathological changes at the maternal-fetal interface could be a cause and/or effect of hypoxia, hyperoxia, oxidative stress and inflammation can cause placental insufficiency and fetal growth retardation. To validate these results, expression of oxidative stress (HIF1 and NRF2), anti-inflammatory (IL-4 and IL-10) and pro-inflammatory (IL-6 and TNFα) markers were studied by semi-quantitative PCR (figure-1). The protein expression of transcription factor NF-κB was evaluated by IHC.

**Figure 1:**
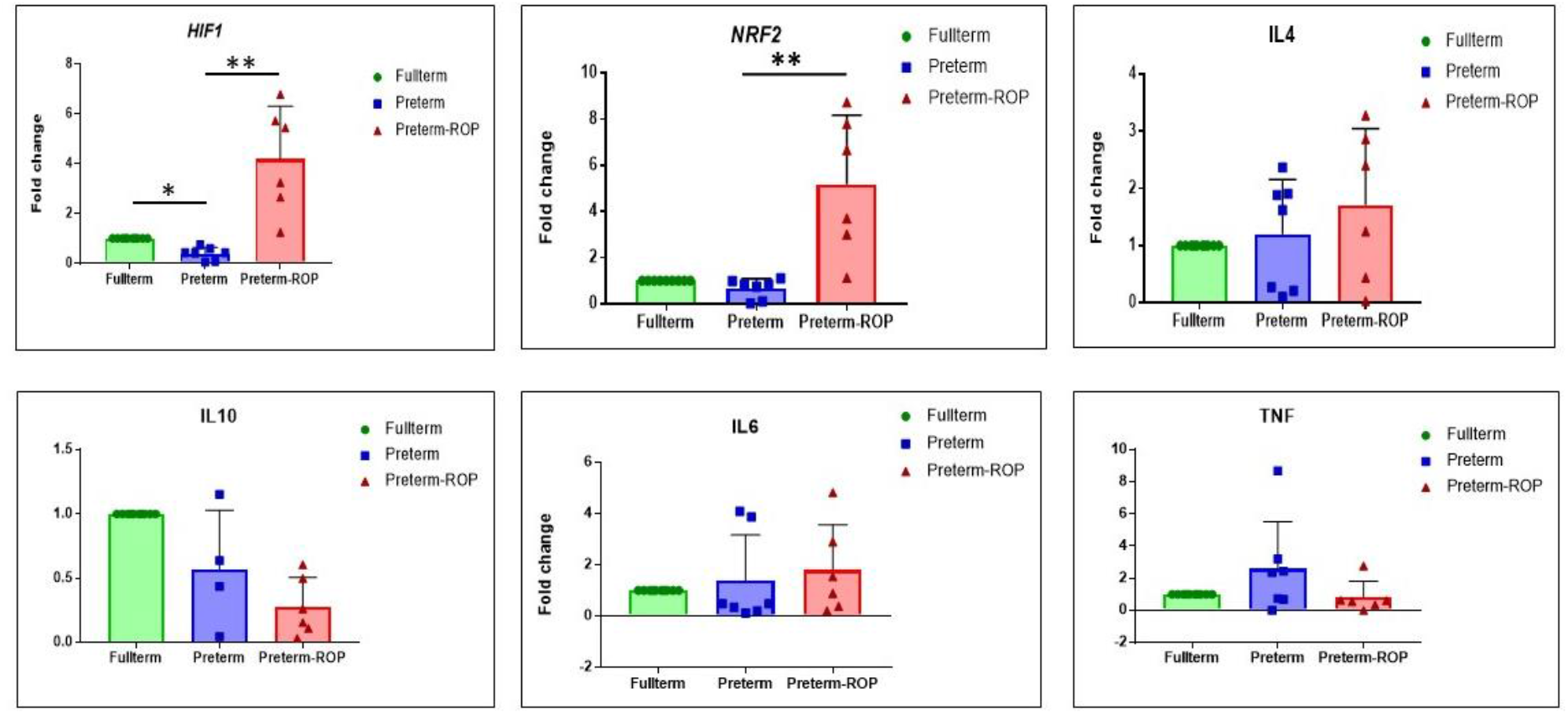
Relative gene expression by qRT-PCR of oxidative stress markers *HIF1* and *NRF2*; anti-inflammatory markers *IL-4* and *IL-10* inflammatory markers *IL-6* and *TNFα* in three studied groups i.e., full term (controls), preterm and preterm with ROP.

### Expression of complement pathway related genes/proteins in placenta at maternal fetal interface

The levels of *C3* gene expression at mRNA level was highest in preterm with ROP but was not significant when compared to other groups. However, the levels were non significantly decreased in preterm without ROP group compared to full-term study group. Same trend was followed when comparison was made for mRNA expression levels of *CFH* gene (figure-2b). But when protein expression was quantified by IHC, the C3 levels were highest in preterm with ROP group followed by preterm without ROP and full-term specifically in the trophoblastic layers of the choronic villi and focally in the fetal blood in capillaries (figure-4). The CFH protein expression was significantly increased in preterm with ROP as compared to preterm without ROP (p≤0.05) and preterm without ROP compared to full-term placentas (figure-3).

**Figure 2:**
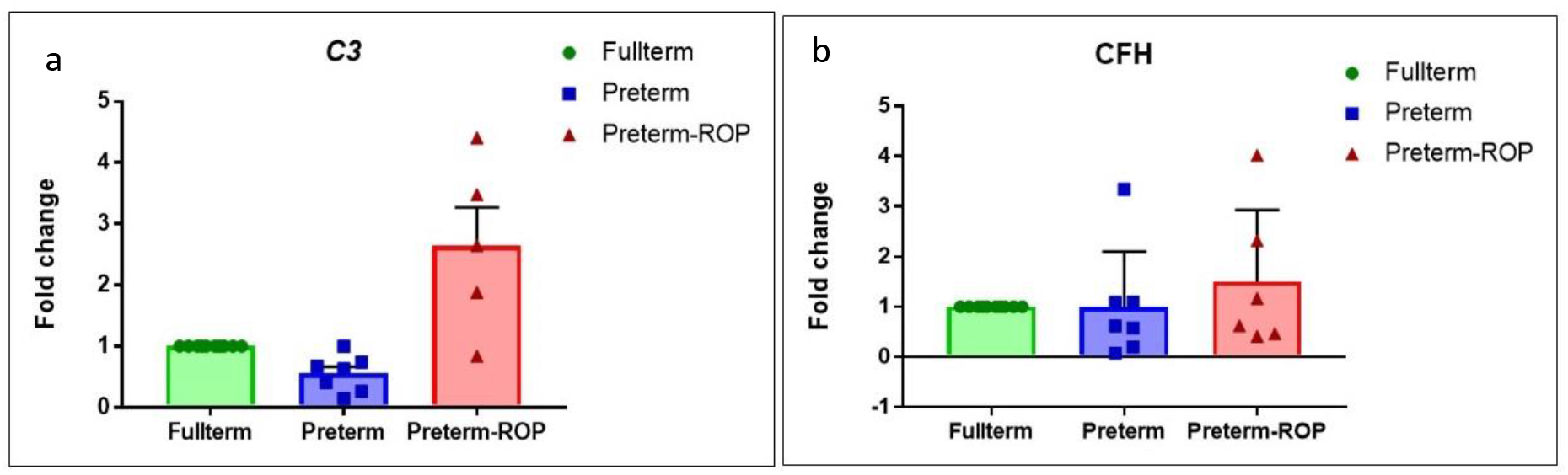
Gene expression of complement system related markers. A) represents comparison of *C3* gene expression; b) represents the comparison of *CFH* gene expression

**Figure 3:**
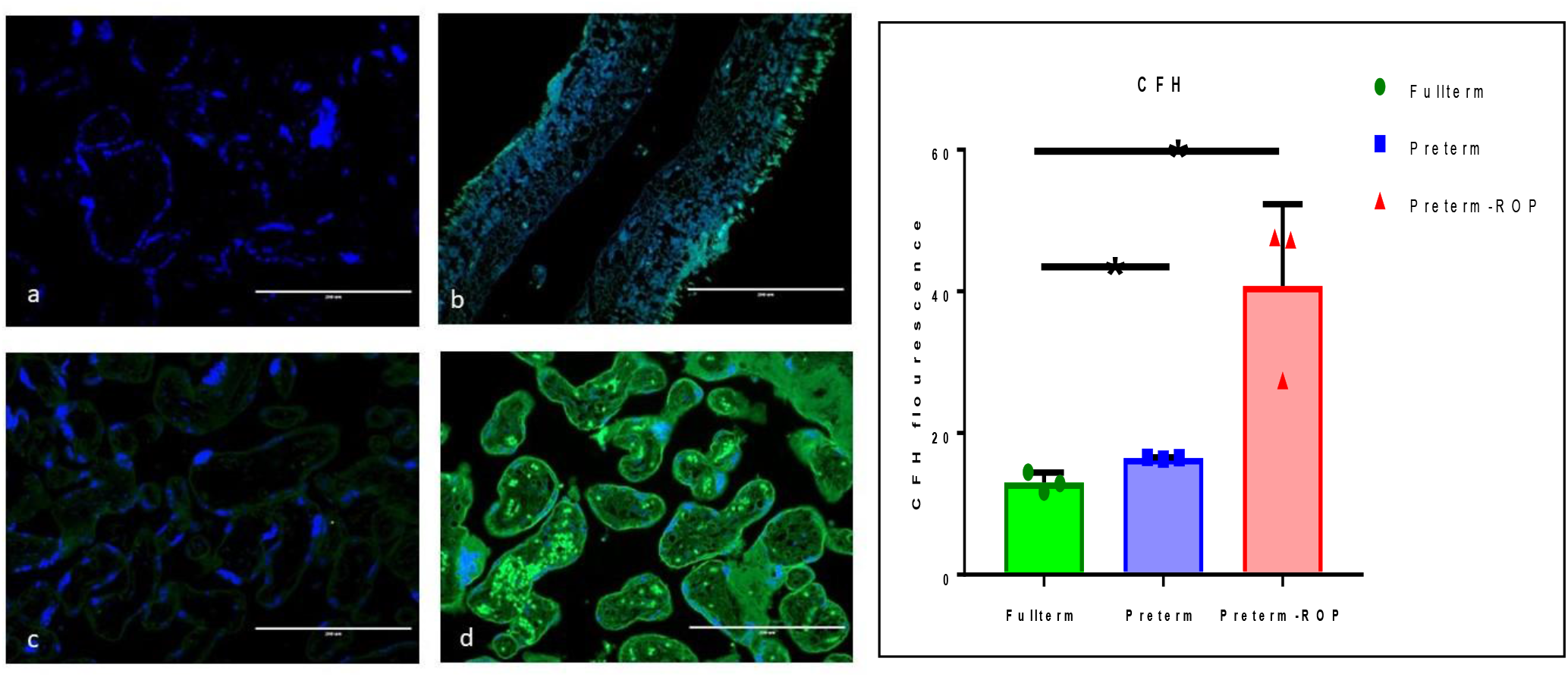
IHC staining with CFH antibody of placental chronic villi from preterm with and without ROP. a) represents no antibody staining for placental tissue; b) positive staining of retinal tissue with CFH antibody; c) preterm without ROP placental chronic villi. d) chronic villi from preterm with ROP placenta showing strong staining with CFH antibody in trophoblastic layers and fetal vessels as compared; graph represents the comparison of CFH protein expression in three groups.

**Figure 4:**
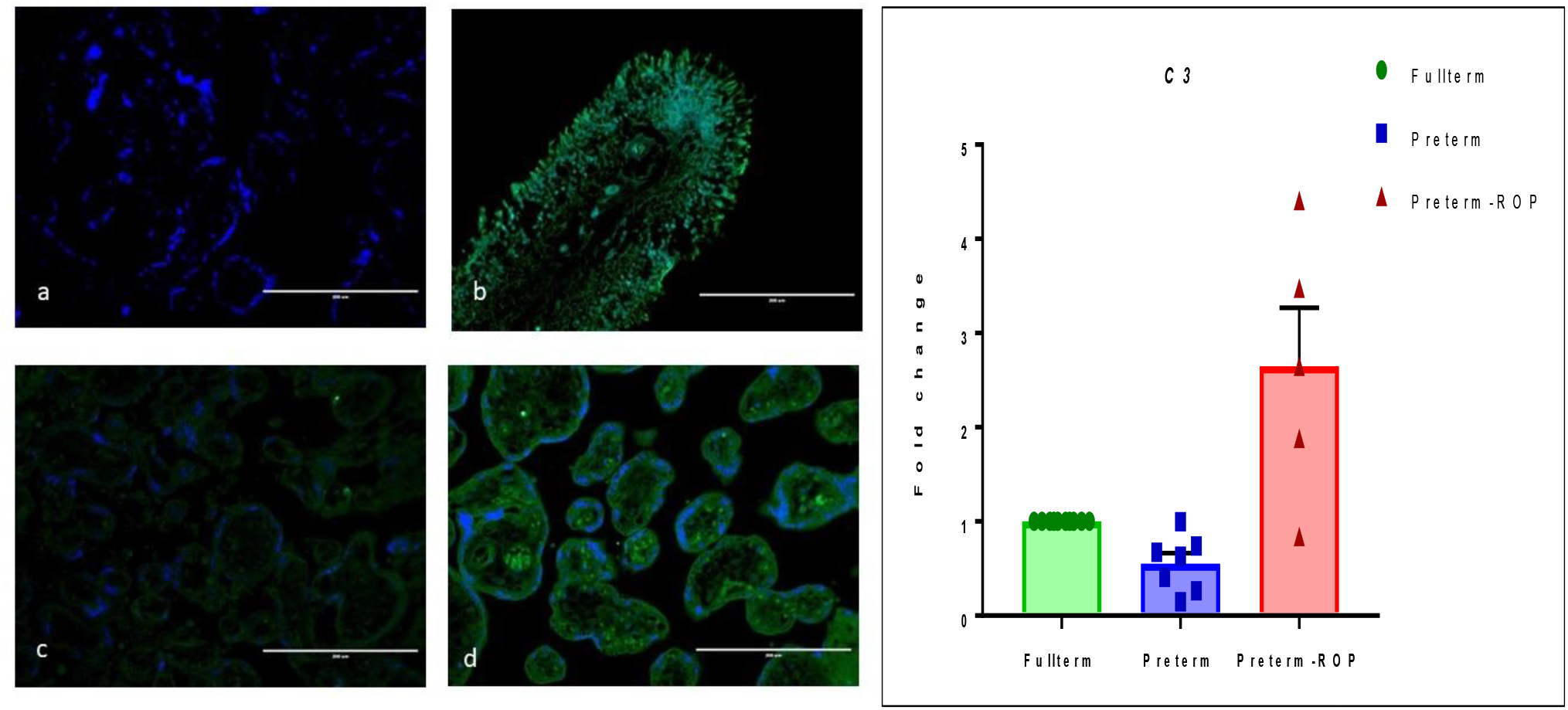
IHC staining with C3 antibody of placental chronic villi from preterm with and without ROP. a) represents no antibody staining for placental tissue; b) positive staining of retinal tissue with C3 antibody; c) preterm without ROP placental chronic villi. d) chronic villi from preterm with ROP placenta showing strong staining with C3 antibody in trophoblastic layers and fetal vessels as compared to preterm without ROP. Graph represents the comparison of C3 protein expression in the three study groups.

### mRNA expression of oxidative stress, inflammatory and anti-inflammatory markers at maternal fetal interface in placenta

When compared the gene expression of oxidation stress related genes i.e., *HIF1* and *NRF2*; *HIF1* expression was significantly higher in the preterm with ROP (p=0.007) compared to preterm without ROP group, while it was significantly decreased in preterm without ROP group compared to full term (p=0.006). *NRF2* gene also followed the same trend as *HIF1* but statistical significance was achieved only when comparison was made between preterm with ROP and Preterm without ROP.

When compared, the expression of anti-inflammatory genes; *IL-4* was non-significantly increased in preterm with ROP compared to preterm without ROP and preterm without ROP compared to full-term. While the expression of *IL-10* was highest in full term followed by preterm without ROP and then preterm with ROP, but this decrease in expression was statistically non-significant. On the other hand, pro-inflammatory marker *IL-6* was highest in preterm with ROP compared to preterm without ROP and full-term but here also the differential expression was not statistically significant. Alternatively, the TNFα levels were non-significantly increased in preterm without ROP group followed by full-term and then preterm with ROP.

But when NF- ΚB protein expression was studied through IHC, there was a significant increase (p=0.022) in the preterm with ROP group compared to preterm without ROP group.

## Discussion

The present case-control study was performed to investigate the impact of maternal, fetal and placental factors on the development of ROP. The preliminary data does show the association of placental histopathological and molecular changes along with maternal and fetal factors towards the disease. There is a clear association between increased monocytes in third trimester, fetal growth retardation, apnoeic spell, ventilation and length of hospital stay and decreased RBC, Hb, PCV at the time of hospital administration, gestational age, birth weight were increasing the risk of the disease. But after multiple linear regression analysis only fetal growth retardation was significantly associated with ROP.

Along with these factors placental weight, diameter was decreased in the preterm with ROP group and significantly associated with the disease when we categorized our study groups into preterm with ROP and preterm without ROP. The previous reports from the literature have also shown the association of reduced placental size with low birth weight and fetal growth retardation (Liu *et al*., 2019; Freedman *et al*., 2019). Specially the surface area of the placenta is known to have a strong association with birth weight and this can be due to reduced number of uteroplacental arteries that supply nutrients to the fetus; reduction in the placental supply to the fetus results in fetal growth retardation (Gaccioli and Lager, 2016). In the present study we have found a strong association of fetal growth retardation with ROP disease. In literature, it has been found that the infants that have prenatal intrauterine growth retardation are at increased risk of developing severe ROP (Chu *et al*., 2020)

Besides the placental size we have also found significant association of vascular changes at the maternal-fetal interface: distal villous hypoplasia, thin elongated villi, Tenny Parker changes, alternating area of small and short hyper mature villi with the disease. Along with these placental changes we have also found rare pathological changes in the placentas from the deliveries where infants developed ROP. These changes are as follows (Figure 5a-5f):

**Figure 5:**
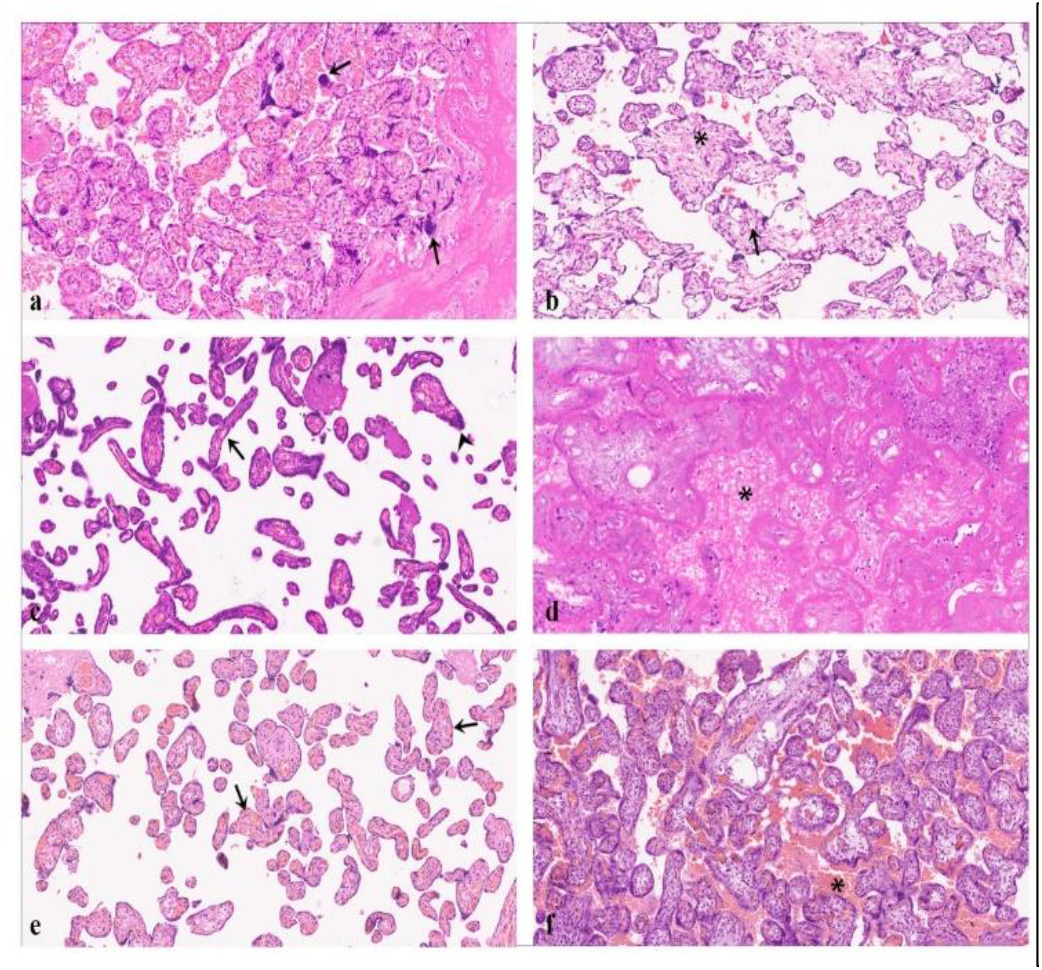
Photomicrographs represents the rare histopathological changes observed in preterm with ROP and without ROP placentas; a) syncytial knotting (Tenny parker changes) observed in majority of full term and preterm without ROP cases (H&E; 10X): arrows represent the syncytial knots; b) Immature villi observed in placenta with GA 34+1 week and baby was scanned as both eyes zone II anterior no plus (H&E; 10X): asterisk represents the immature villous and arrow represents the deep fetal capillary; c) Distal villous hypoplasia observed in placenta with GA 29+3 weeks and baby was scanned as both eyes zone II, avascular retina (H&E; 10X): arrows represent the slender shaped chronic villi and arrow heads represent the syncytial knots; d) Ghost villi observed in placenta with GA 28+2 weeks and baby was scanned as both eyes zone II, immature retina (H&E; 10X): asterisk represents the accumulation of debris from dead cells into the intervillous space; e) chorangiosis observed in placenta with GA 29+5 weeks and baby was scanned as both eyes zone II, no plus (H&E; 10X): arrows represent the chronic villi with more than 10 fetal capillaries; f) haemorrhage observed in placenta with GA 28+1 weeks and baby died after five days of life.

### Syncytial knotting (Tenny-Parker changes)

when syncytial nuclei aggregate at the surface of terminal villi, they are known as syncytial knots. The condition is associated with uteroplacental malperfusion and conditions linked with hyperoxia, hypoxia and reactive oxygen species. The syncytial knots increase significantly in placentas from pre-eclamptic and fetal growth restriction (FGR) pregnancies (Heazell *et al*., 2007). Both preeclampsia (Shulman *et al*., 2017) and fetal growth retardation (Allegaert *et al*., 2003) are strongly associated with ROP.

### Immature Villi

The immature villi lead to reduction in the fetoplacental weight ratio which is considered as a measure of placental efficiency. The immature villi lack specialized vasculo-syncytial membranes and fetal capillaries are deep seated in these immature villi that results into increased diffusion distance and time for supplements supply to fetus thus resulting to intrauterine growth retardation (Stallmach and Hebisch, 2004).

### Distal villous hypoplasia / terminal villous deficiency

In distal villous hypoplasia, villous tree is poorly evolved, shape of the villi is elongated and slender and hence can cause increased intervillous space. This increase in intervillous space can further result into intrauterine growth restriction with unfavorable fetal sustainability and can cause lifelong neurovascular developmental and cardiovascular health issues (Fitzgerald *et al*., 2021).

### Ghost villi

Chorionic villi in a placental infarction that may or may not be surrounded by inflammatory cells. Ghost villi are results of focal decrease in the placental blood supply leading to ischemic necrosis in the former (Beebe *et al*., 1996).

### Chorangiosis

Extreme terminal villous hyperplasia known as chorangiosis is defined by Altshuler in 1984 as, “chorangiosis is characterized by more than 10 capillaries, in more than 10 chronic villi in several areas of placenta” (Altshuler, 1984). The prevalence of chorangiosis is increased in the presence of other acute and/or chronic hypoxia lesions. The hypervascularity is formed due to low oxygen in the villous capillaries (Suzuki *et al*., 2009).

### Intervillous haemorrhage

Hemorrhage within the villous stroma occurs in early stages of an acute hypoxic event. Most cases are observed in association with abruption or recent infarcts, and therefore, this phenomenon is observed more often in disorders underlying maternal vascular malperfusion (Meir and Ariel, 2019).

In the gestational environment these histopathological changes can cause or effect levels of inflammation (Goines *et al*., 2011), hyperoxia and hypoxia (Burstyn *et al*., 2011) which are considered to play an important role in neurodeveleopment disablities (walker et al., 2015). when trophoblastic layers are exposed to these stresses, increased levels of sVEGF-R1 is released into maternal circulation (Maynard *et al*., 2003) this prevents trophoblastic differentiation and hence abnormal placentation (Charnock-Jones *et al*., 2000). The increased levels of antiangiogenic factors in maternal circulation during different development stages can delay the vascular development of the fetal organs including retinal vascularization that can also leads to the development of ROP in postnatal life (Shulman *et al*., 2017)) Along with this, increased VEGF-R1 is released due to stressed fetal layers at maternal-fetal interface is also a determinant of decreased levels of vascular endothelial growth factor (VEGF), an important feature of phase-I of ROP (Alshaikh *et al*., 2017). Thus, the observed placental histopathological changes and their *in-utero* consequences during fetal development align with the phase-I manifestations of ROP.

The increased expression of complement system related genes *C3* and *CFH* at maternal fetal interface indicates the possible activation of alternative complement pathway in placentas from the infants that developed ROP in postnatal life. The fetal syncytiotrophoblast the outer most layer of chronic villi is the only layer that is in direct contact with the maternal blood. So, complement activation at this cellular layer should be carefully controlled to avoid the harm to fetus from maternal immune system (Regal *et al*., 2015). The increased expression of these proteins at maternal-fetal interface can further triger the inflammatory response. Uncontrolled inflammation leads to unusal pathologies related to placetal villi damage characterized by apoptosis and can alter the normal functioning of trophoblastic layers at terminal chronic villi (Genbacev et al., 1999; Rampersad *et al*., 2008) that can add to impaired development of utero-placental system and blood supply to fetus (Regal *et al*., 2015) and can also underwrite to pathologies like preeclampsia (Huppertz, 2018). Increased expression of maternal complement proteins is known to be associated with insufficient trophoblast invasion and arterial remodelling (Regal *et al*., 2017)) leading to reduced blood flow and conditions like hypoxia and increased chances of fetal growth retardation. These small for age foetuses are at increased risk of neurodevelopmental disorders after birth (Colson *et al*., 2021). In normal conditions, the complement activation is downregulated in the preterm infants due to immature immune system but we have found increased levels of these factors in preterm with ROP placentas indicating their possible role in ROP pathogenesis. In a previous study from our lab, we have found a strong association of complement system related gene variants with ROP and also increased expression of proteins in the extra-cellular matrix and vitreous of infants with ROP (Rathi et al., 2017). The levels of complement system related proteins are low in preterm babies because of immature immune system (Grumach *et al*., 2014) the present study results confirms that the activation and accumulation of abnormal immune complement in the eyes of these babies that develop ROP can be a consequence of in-utero changes in ROP.

Together with histopathological hypoxic lesions, significant increase in the *HIF1* and *NRF2* gene expression in preterm with ROP placentas compared to preterm without ROP indicates the presence of oxidative stress at maternal-fetal interface. The tissue oxygen levels in hypoxic conditions is mediated by *HIF1*. In hypoxic conditions HIF1 conciliates oxygen levels by inducing the expression of erythropoietin (EPO) which inturn increases RBC production and their oxygen carrying capacity (Goldberg *et al*., 1988). The HIF1 is heterodimer of two subunits HIF1-α and HIF1-β and its activity is mainly controlled by HIF1-α subunit in oxygen dependent manner. It also controls the process of angiogenesis in hypoxic conditions. Angiogenesis is a complex cascade of events involving multiple genes and proteins. In hypoxic state, many of these genes gets upregulate to fulfil the tissue oxygen demand (Levy *et al*., 1995; Forsythe *et al*., 1996; Bunn and Poyton, 1996). One of the mechanisms by which HIF-1 regulates angiogenesis is by inducing the VEGF expression for upregulation of endothelial cell proliferation (Josko *et al*., 2000). Due to its control over regulation of oxygen supply, HIF1 plays important role in ocular diseases including ROP. Retina is a light sensitive tissue and requires continuous oxygen supply to full fill energy demands for proper functioning of specially rod and cone cells which is provided by choroidal and retinal circulation and both are important to maintain the retinal physiology and lack of oxygen can lead to visual impairments (Pournaras *et al*., 2008). The retinal vasculature is incomplete in the preterm infants and they also need supplementary oxygen after birth because of immature lungs. The retinal vasculature takes total 40 weeks to complete, in preterm infants retinal vasculature is not fully formed and hence require supplement oxygen after birth to full-fill the oxygen demand for metabolic activities and energy production. The sudden increase in the oxygen levels leads to hyperoxia and vaso-obliteration in the retina (phase-I) of the disease followed by hypoxic stress in phase-II when the infants return to room air. This activates the HIF1-α that activates angiogenic factors leading to neovascularization in retina to overcome oxygen insufficiency. This excessive vascularization can lead to retinal detachment and hence vision loss (Chen *et al*., 2007; Sapieha *et al*., 2010). Hence, levels of HIF1-α are important as oxygen tension which is one of the major contributors to ROP is regulated by the former. In company with the *HIF1-α*, *NRF-2* is also required to maintain the oxygen levels as HIF1-α regulates hypoxia, *NRF-2* regulates oxidative stress but in case of intermittent hypoxia both *HIF1-α* and *NRF2* pathways were induced (Malec *et al*., 2010. Both angiotensin II receptor I signalling in preeclamptic patients results in increased ROS production (Nguyen *et al*., 2013) along with the other factors like TNFα. But increased NRF2 levels results in the decreased expression of TNFα, VGFR1 and other factors (Zhang *et al*., 2021). Our results are in concordance with the observations from Zhang and co-workers (2021) as there was non-significant increase in NRF2 levels at the maternal-fetal interface in placentas from infants that develop ROP compared to preterm placentas without ROP, while TNFα levels were decreased.

There were no significant changes in the levels of expression of pro-inflammatory (IL-6 and TNFα) and anti-inflammatory (IL-4 and IL10) markers at the maternal fetal interface. A report by Woo *et al*., 2013 had observed a significant association of Apgar score with ROP but could not find any association of levels of inflammatory cytokines with ROP (Woo *et al*., 2013). However, in a study that had evaluated association of 10 cord blood serum cytokines with ROP, significant association of IL-7 monocyte chemotactic protein-1, macrophage inflammatory protein 1 alpha, and macrophage inflammatory protein 1 beta was seen (Yu *et al*., 2014). Such inconsistencies in results can be due to smaller sample sizes or due to ethnic differences across the study groups. In another study, by Park and co-workers in 2019 found a strong association of increased IL-6 and C5a levels with the risk of developing the ROP. IL-6 is a potent pro-inflammatory cytokine and a well-established risk factor for ROP (Hartnett, 2015) as well as for several fetal inflammatory complications (Goepfert *et al*., 2004; Su *et al*., 2014). Although we have not found any significant change in the pro- and anti-inflammatory markers in ROP and no-ROP placental samples but there is requirement of further validation on a larger sample size.

This is a very preliminary data that indicates the involvement of several prenatal factors that lay down the foundation for the postnatal pathological changes leading to the neovascularization in the retina of the eye and further can result to permanent vision loss in the preterm infants. The results of the present study can be used in future to develop diagnostic test and comprehensive management of the disease.

## Conclusion

To our knowledge the relationship between the placental histopathological and molecular changes at the maternal-fetal interface and ROP has not been studied previously so our study is unique in this respect. The association of these changes with the disease supports our hypothesis that there are in-utero pathological changes in the placenta that predisposed the preterm infants to increased risk of ROP development. But the results are needed to be further validated in the larger cohort.

